# Enhancing backcross programs through increased recombination

**DOI:** 10.1101/2020.05.05.078287

**Authors:** Elise Tourrette, Matthieu Falque, Olivier C. Martin

## Abstract

Introgression of a QTL by successive backcrosses is a strategy that can be used to improve elite lines (recurrent parent) by bringing in alleles from exotic material (donor parent). In the absence of selection, the proportion of the donor genome decreases by half at each generation. However, since one selects for the donor allele at the QTL, the elimination of the donor genome in the neighborhood of that QTL will be much slower (linkage drag). Using markers to monitor the genome around the QTL and in the background can accelerate the return to the recurrent parent genome. The success of an introgression will partly depend on the occurrence of crossovers at favorable positions. However, the number of crossovers per generation is limited and their distribution along the genome is heterogeneous. Recently, techniques have been developed to modify these two aspects of recombination. Here, we assess, by simulation, their effect on the efficiency of introgression programs by studying the reduction of the linkage drag and the recovery of the recurrent genome. The selection schemes we simulate begin by two generations of foreground selection and continue with one or more generations of background selection. Our results show that when the QTL is in a region that was initially lacking crossovers, increasing the recombination rate can decrease the linkage drag nearly ten-fold after the foreground selection and improves the return to the recurrent parent. However, if the QTL is in a region already rich in crossovers then increasing recombination proves to be detrimental.

**Key message:** In breeding programs, recombination is essential for introgression, but introducing more crossovers is beneficial only when the target is in a cold region, otherwise it is detrimental.

## INTRODUCTION

Breeding schemes based on recurrent backcrossing introgress an allele from a donor parent at a target locus into the genetic background of a recurrent parent. Beyond the need to maintain the donor allele of the target locus in the progenies, such a scheme has two aims: (1) reduction of the segment size from the donor parent around the target locus and (2) recovery of the recurrent parent genomic background. These objectives are achieved by doing multiple generations of backcrosses between the offspring of the previous cross and the recurrent parent (Hospital 2005). Backcross has multiple uses in genetics and breeding, ranging from validating a putative allelic effect to genetically improving agriculturally important species. The method aims to transfer a favorable allele of one or multiple loci previously in a poor genetic background into a better one. Examples include the introgression of a resistance gene from a non-elite genotype like a landrace, the introgression of a transgene into a reference line (if there are multiple loci, the process is called gene pyramiding, Servin *et al.* 2004), unraveling the genetic architecture of a quantitative trait, testing for additivity of the effects of QTLs (Quantitative Trait Locus), or increasing the precision in QTL mapping (Hospital 2005, Tanksley and Nelson 1996)]. In the absence of any selection, the proportion of the donor genome will be halved at each generation. However, since the target locus is linked to its neighboring loci, the selection at the target locus will generally also select for the donor genome on those loci. As a result, the proportion of the donor genome will decrease less for the chromosome carrying the target locus (carrier chromosome). This is the so called linkage drag problem (Hanson 1959, Stam and Zeven 1981, Naveira and Barbadilla 1992). It is possible to accelerate the return to the recurrent genome and to reduce the linkage drag by exploiting markers both in the genetic background and close to the target locus (Young and Tanksley 1981, Visscher *et al.* 1996, Frisch *et al.* 1999b, Hospital 2001).

The reduction of the linkage drag around the target locus depends on recombination: in the absence of recombination events the linkage drag would extend to the whole carrier chromosome. Since there is almost always some recombination, one can use multiple generations to accumulate recombination events (that is true even in the presence of crossover interference which prevents close-by crossovers from being formed during the same meiosis; indeed crossovers are independent if one considers different meioses and thus different generations). Hence, it is possible to obtain crossovers closely flanking the target locus (on both sides) if sufficiently many generations are used, even in the presence of crossover interference.

There are a number of methodological studies on backcross programs that investigate the role of population size or location of markers. For instance several authors considered how to optimize the positions of a limited number of markers flanking the target locus (Hospital and Charcosset 1997, Visscher *et al.* 1996). In particular,Hospital *et al.* 1992 and Frisch *et al.* 1999a concluded that the bigger the population, the closer the markers should be to the target locus. Frisch and Melchinger (2005) theoretically studied the proportion of donor genome at one generation depending on the genetic map and the markers’ genotypes. Rodolphe *et al.* (2008) were interested in the parental composition of the chromosomes, that is they considered the statistics of the mosaic chromosome structure as a function of generations, providing the distribution of sizes of the chromosomal blocks coming from the donor or the recurrent parent; they did so assuming two models of recombination: recombination without interference and recombination with complete interference, *i.e.* one crossover per chromosome at each generation.

Recombination events arising during a backcross program will influence the degree to which one can return to the recurrent genome. Thus, in the present work we ask whether increasing recombination might speed up introgression schemes.

A few experimental techniques have been developed to increase recombination rates or modify recombination landscapes. One method knocks out anti-crossover genes (Fernandes *et al.* 2018 in *Arabidopsis thaliana* and Mieulet *et al.* 2018 in pea, rice and tomato). Another method is based on modifying the ploidy levels in *Brassica rapa* (Pelé *et al.* 2017). Both of those methods lead to many-fold increases in crossover rates and interestingly, the second one also affects the recombination profiles by adding crossovers to the crossover-poor regions. More modest increases (up to two-fold) have been obtained through over-expressing pro-crossover genes (Ziolkowski *et al.* 2017 and Serra *et al.* 2018 in *A. thaliana*). A completely different approach consists in targeting crossovers at some particular genomic locations by transgenesis (Choi 2017) which increases recombination rates a lot but only very locally. Beyond these methods that manipulate recombination by biotechnological means, it is possible to exploit natural variations in recombination rates. These arise as differences between male and female meiosis (for instance as shown by Giraut *et al.* 2011 in *A. thaliana* and Phillips *et al.* 2015 in barley), as differences due to genetic backgrounds (Salomé *et al.* 2012 and Bauer *et al.* 2013) or as responses to different environmental conditions (Lloyd *et al.*, 2017, Modliszewski and Copenhaver 2017). Blary and Jenczewski (2019) provide a review of all these cases. In *Arabidopsis thaliana*, one of the double-mutants of anti-crossovers genes increased the recombination rate 7.8-fold (Fernandes *et al.* 2018) *via* the production of additional non-interfering crossovers. This decrease of interference increases the probability that, in a single meiosis, two crossovers occur close to each other, for our purposes potentially on both sides of the target locus, in backcross schemes. Such a property may speed up the linkage drag reduction since one could bypass the need to select for a crossover on each side of the target locus in different generations (what is normally done in two generations could perhaps be done in one).

In the present work, we use an *in silico* approach to determine the impact of increasing recombination and modifying recombination landscapes in backcross programs, focusing on the objective of recovering as much as possible of the recurrent genome. Specifically, we simulate backcross programs under normal and increased/modified recombination schemes as given by the experimentally observed increases in the first two approaches. These schemes respectively use (1) mutants of anti-crossover genes and (2) modifications of the ploidy level; hereafter we shall refer to these two approaches as HR (for “HyperRecombinant”) and Boost (for boosted recombination). We use as our main focus a “standard” backcross program in which after BC3 (third generation of backcross) there is one generation of selfing (program BC3S1) but we also consider alternative programs. Based on our investigation of how successful such backcross programs are depending on various choices such as the population sizes or the position of the target locus, we find that modifying recombination rate is generally quite advantageous if it changes the region containing the target locus from being cold to warm (or even hot) with respect to recombination.

## MATERIALS AND METHODS

Using modeling and simulations, we investigated the effects of increasing recombination or modifying recombination landscapes on a program introgressing the allele of a donor parent at a target locus into a recurrent parent by successive backcrosses. Our computer codes were written in the programming language R (R Core Team 2018), both for producing individual-based simulations of forward-in-time backcross breeding schemes and for all the associated analyses.

### Recombination landscapes and simulation of crossover formation

Different levels of recombination were compared (see Fig. 1, Fig. S1 and Fig. S2 for the different recombination landscapes). The WT situation corresponds to the normal level of recombination while Boost and HR correspond to the increased levels of recombination as produced by the two methods reviewed in the introduction. Under Boost, the modified recombination landscape has an increased recombination rate in the pericentromeric regions which are normally almost void of crossovers. This landscape was experimentally measured in the context of plants having a non standard karyotype, specifically some chromosomes were present as pairs (diploid state) while others were present as singletons (haploid state). It is believed that this mixed state perturbs regulatory processes and thus allow for a large increase in crossover number. For our *in silico* study, we assume that this modified landscape will arise in all the successive generations of the backcross program. In practice, that may mean that only a subset of the progenies (individuals having sufficiently many chromosomes in the haploid state) can contribute to the program. Under HR, the recombination rate is globally increased without significant changes in the shape of the recombination profile, the pericentromeric regions remaining poor in crossovers. The recombination landscapes of WT and Boost were taken from published results in *Brassica rapa* (Pelé *et al.* 2017) while the landscape of HR was simulated based on published results in *Oryza sativa* (Mieulet *et al.* 2018), as it was not available in *B. rapa*. Specifically, we simulated the HR landscape to reproduce the absence of any change in *shape* of the landscape compared to the WT one. To do so, the WT landscape was multiplied by a constant. The constant, dependent on the chromosome of *B. rapa*, was chosen to get the same chromosome-wide increase of the recombination as for Boost; it was thus set to *L_G_Boost_*/*L_G_WT_*, *L_G_* denoting the genetic lengths (see Table 1 for the genetic lengths under normal and increased recombination). Thus, by considering Boost and HR, we were able to observe the effects of changing or not changing the shape of the recombination profiles when increasing the recombination rate.

**Figure 1.**
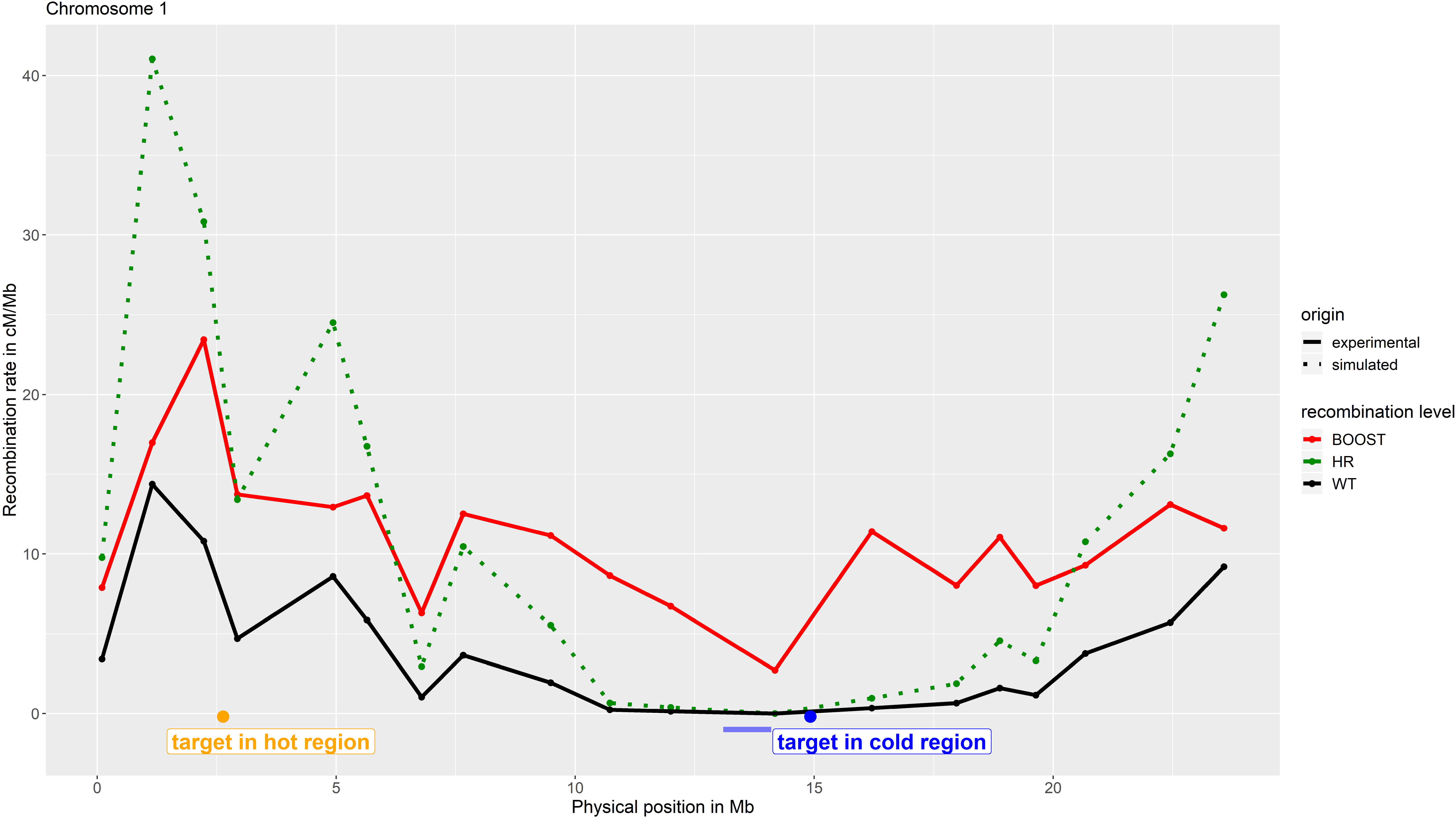
Recombination landscape of chromosome 1 of *Brassica rapa*. The WT landscape is represented in black while the HR is in green and Boost is in red. The solid lines represent the profiles obtained from experimental data (WT and Boost, data from Pelé *et al.* 2017) while the dotted line is the simulated profile (HR). The yellow and blue dots represent the positions of the QTL in a hot and cold region, respectively (positions taken from Kumar *et al.* 2018). The centromere position is represented by the blue bar (Mason *et al.* 2016).

**Table 1.** Genetic lengths, in centiMorgans, for the ten chromosomes of *B. rapa*, under normal and increased (HR and Boost) recombination (taken from Pelé *et al.* 2017)

| Chromosome | *B. rapa* WT | *B. rapa* Boost and HR |
| --- | --- | --- |
| A01 | 92.9 | 265.2 |
| A02 | 78 | 260.7 |
| A03 | 102.9 | 396.3 |
| A04 | 56.2 | 206.5 |
| A05 | 95.4 | 263.3 |
| A06 | 105.7 | 300.9 |
| A07 | 74.1 | 318.3 |
| A08 | 56.8 | 225.8 |
| A09 | 103.9 | 394.7 |
| A10 | 55.8 | 143.1 |
| TOTAL | 821.7 | 2774.8 |

Since we had the physical positions of the target locus and of the markers, it was necessary to calculate their genetic positions in each case (WT, Boost and HR). Using the different recombination landscapes, we interpolated the relationships between genetic and physical positions using splines (R function smooth.spline with spar = 0.1 to remove noise in the data). In our comparisons of these three cases, the genetic positions of the markers changed while their physical positions remained constant.

Meiotic recombination was simulated by generating crossovers in genetic space. The number of crossovers was drawn from a Poisson distribution, following the model of Haldane (McPeek and Speed 1995) that supposes crossover formation without genetic interference. Hence, the genetic distance between two adjacent crossovers could be drawn from an exponential distribution, with an average of one crossover per Morgan.

The markers were regularly spaced along the chromosomes (in physical coordinates): there was one marker every 250 kb (4 per Mb) in the genome-wide background and 100 times more in a 2 Mb interval around the target locus (one every 2.5 kb) in order to be more precise during the foreground selection.

The target locus was located either in a cold region (at position 15 Mb) or in a hot region (at position 2 Mb) on chromosome 1 (these physical positions for the QTLs, see Fig. 1, are taken from Kumar *et al.* (2018)).

### Backcross breeding scheme

The backcross scheme (Fig. 2) consisted in a first cross between a donor parent, which has the wanted allele at the target locus, and a recurrent parent, having the desired genetic background. Following this cross producing an F1, we simulated a succession of backcrosses between one offspring of the previous cross and the recurrent parent. Across successive generations, we aim to recover as much as possible the recurrent background while keeping the donor allele at the target locus.

**Figure 2.**
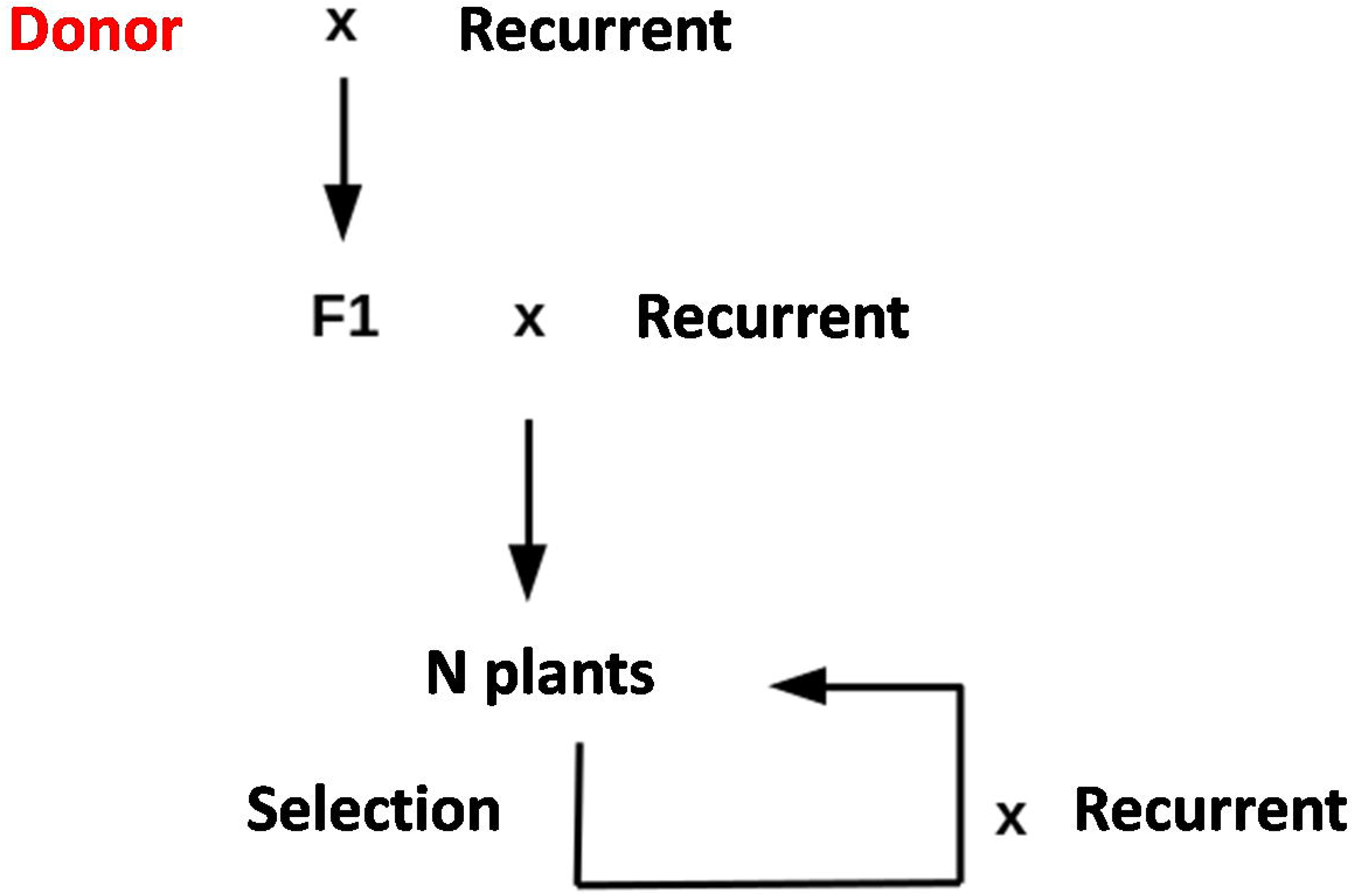
Scheme of a general introgression breeding program using successive backcrosses. The scheme can end by one generation of selfing (this generation is noted as BCnS1).

For the reference situation, presented in detail in the main part of this paper, we use three or four generations of backcross followed by one generation of selfing. For better theoretical understanding, an additional situation of eight generations of backcross was also studied (presented in the Supplemental Material). At each generation, the population consisted of 400 plants (additional situations treated in the Supplemental Material use instead 200 or 1000 plants). At each generation, only one plant was selected to be backcrossed to the recurrent parent at the next generation.

The selection criteria applied to select that single plant occurred in two phases. First we pre-selected the plants carrying the donor allele at the target locus (around half of the population was thus kept in this phase). For the second phase, the selection criterion depended on the generation considered. In BC1 and BC2 (first and second generation of backcross after the initial cross between the donor and the recurrent), the plant with the closest crossover to the target locus was kept (in BC2 the side for the crossover was the opposite one from that obtained in BC1) in order to reduce the linkage drag (foreground selection). In later generations, we determined for each individual its proportion of donor genome, genome-wide, and the plant with the lowest proportion was kept. This reduction of the donor genome helps to recover the recurrent genome (background selection). To take into account the fact that the best plant may be unavailable (death, sterility), we also looked at the effect of keeping the second best plant instead of the best one.

### Analyses

Two points are important for the success of a backcross program: the reduction of the linkage drag around the target locus and the recovery of the recurrent genome in the genetic background. Hence, in order to assess the effect of increasing recombination on a backcross scheme, we quantified both of these aspects. The linkage drag was evaluated using the length, in Mb, of the donor segment around the target locus. The recovery of the recurrent genome, as for the background selection, was measured using the proportion of donor genome, genome-wide (specifically, the weighted proportion of markers carrying the donor allele, each marker being weighted according to the size of the interval it represents and the number – 0, 1 or 2 – of copies of the donor allele it includes). To further assess the speed of the return to the recurrent genome in the genetic background, we calculated the proportion of donor genome that was due to the linkage drag, *i.e.* that was due solely to the donor segment around the target locus (weighted number of markers in the linkage drag segment divided by the total weighted number of markers as above).

To take into account the stochasticity of the recombination process and thus of the backcross selection program, 500 replicates were generated for each situation and we calculated the mean over the different replicates as well as the 95% confidence interval on the mean (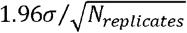, σ being the standard deviation of the considered observable).

## RESULTS

### Effect of increased recombination on the linkage drag

The linkage drag is monitored by the size, in Mb, of the donor segment around the target locus. Let us consider first the most interesting situation which is when the target locus is in a cold region. Fig. 3 shows typical segments while the corresponding mean sizes as a function of generation are shown in Fig. 4.A. In BC1 and BC2, the linkage drag is strongly decreased because the selection focuses on its reduction by keeping the plants with the crossovers closest to the target locus. This decrease is much stronger under Boost than for the WT: in BC1, the mean segment length is 15 Mb for WT and 7.9 Mb for Boost; in BC2, the mean segment length is 5.3 Mb for WT and 0.52Mb for Boost. For the later generations, since the selection focuses on the genome-wide recovery of the recurrent genome, the segment around the target locus hardly decreases with generations. For instance at BC3S1 the mean size of the donor segment at the target locus is 5.2 Mb under WT and 0.51 Mb under Boost. Adding one more generation (up to BC4S1) affects very little those numbers: the mean sizes are 4.3 Mb under WT and 0.49 Mb under Boost. Thus the ratio of sizes under WT and Boost is not sensitive to the addition of an extra generation.

**Figure 3.**
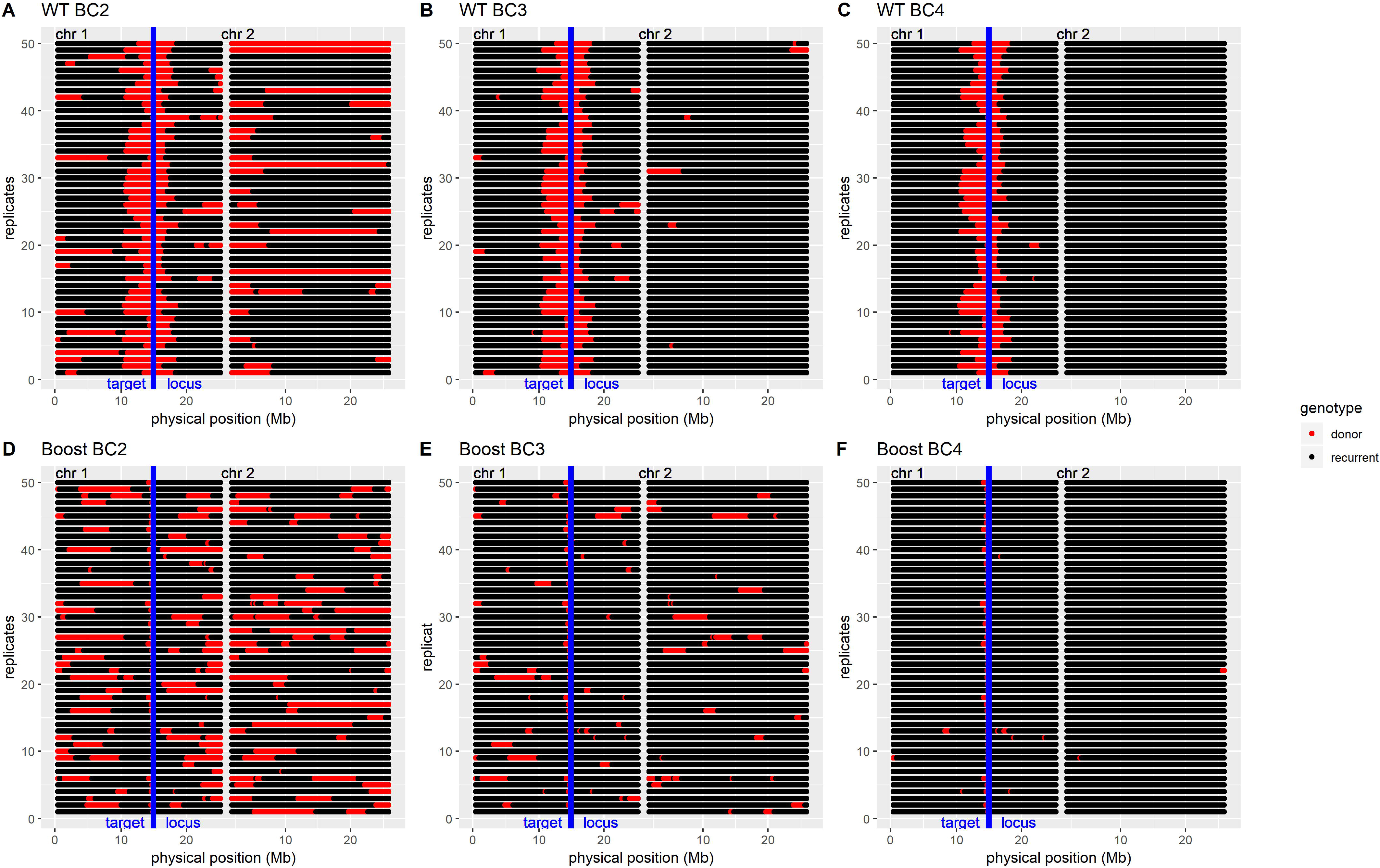
Graphical genotypes (genotypes at markers for 50 random replicates *vs* the physical position) for the generations BC2 (A and D), BC3 (B and E) and BC4 (C and F) under WT (top) and Boost (bottom) recombination, for chromosomes 1 and 2 of *B.rapa*. The markers that are heterozygous (donor/recurrent) are represented in red while those that are homozygous for the recurrent allele are in black. The position of the target locus, on chromosome 1, is represented by a blue line.

**Figure 4.**
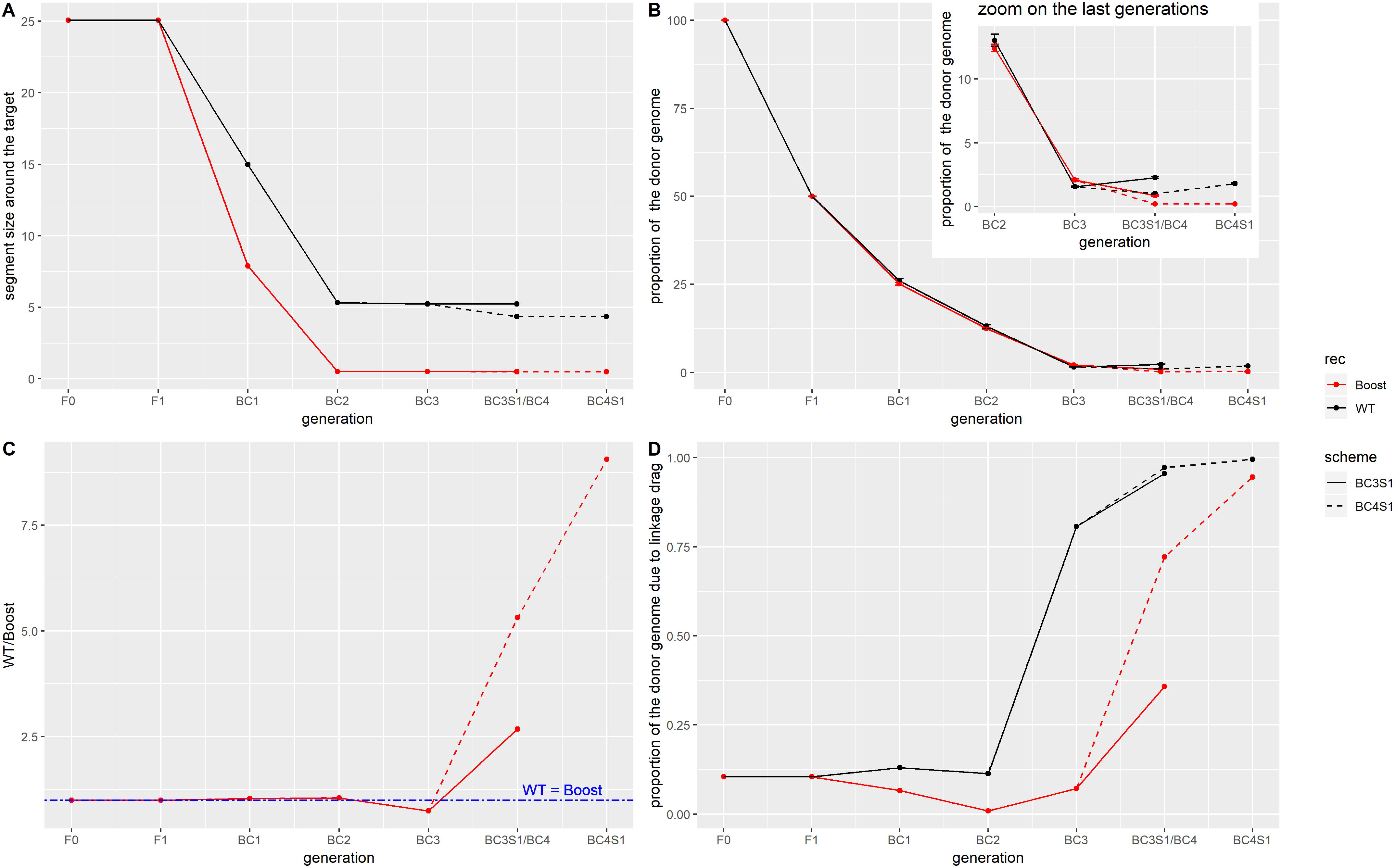
(A) Mean size of the heterozygous segment around the target locus in a cold recombinant region, in Mb, as a function of the generations, in *B.rapa*. (B) Mean proportion of the donor genome, in percentage, as a function of the generations. The insert represents a zoom on the last generations, from BC2 onwards. (C) Ratio of the mean proportion of donor genome under WT over the proportion under Boost as a function of the generations. (D) Mean proportion of the remaining donor genome that is due to the linkage drag, calculated as the part of the remaining donor genome that is due to the heterozygous segment around the target locus, as a function of the generations. The measures under WT are represented in black while those under Boosted recombination are in red. Two selection schemes are shown: up to BC3S1, in solid lines, and up to BC4S1, in dashed lines. The generations BC3S1 and BC4 have the same position on the x axis. In the situations represented in this figure, the target locus is in a cold region, and there are 400 plants per generation. The best individual is kept at each generation, following the selection criterion appropriate for each generation (BC1 and BC2: foreground selection, and thereafter background selection).

Overall, having very little recombination around the target locus is detrimental for the selection program, even though there are 400 plants: at the end of the BC2 generation, the donor segment around the target locus is around ten times bigger for WT compared to Boost, and this is due to the fact that Boost has significantly increased recombination around the target locus (Fig. 1).

The same trend is seen for other population sizes (Fig. S3.A). Increasing the population size decreases the linkage drag, both in WT and Boost: in BC3S1, for a population of 200 individuals, the mean segment size is 6.6 Mb for WT and 0.82 for Boost, while for a population of 1000 individuals, it is 3.5 for WT and 0.24 for Boost. If further generations are added (Fig. S4.A), we observe a progressive decrease of the linkage drag. By going up to BC8, although the selection from BC3 to BC8 is on the genome-wide proportion of donor genome, it also acts on the donor segment containing the target locus. For instance in BC8, we obtain a mean donor segment size of 2.7 Mb for WT and 0.17 Mb for Boost, and this decrease can be explained by the fact that most of the remaining donor genome is due to the linkage drag (*i.e.,* is from the donor segment around the target locus, Fig. 4.D: from BC5 onward, more than 90% of the donor genome is due to the linkage drag). Thus, selecting against the donor genome is similar to selecting to decrease the linkage drag.

Consider now the situation of a target locus belonging to a hot region (Fig. S5.A). Recombination already occurs there frequently in WT. The gain obtained from using Boost is then a lot smaller: for the hot region chosen (Fig. 1), the mean segment size is 0.07 Mb for WT and 0.03 Mb for Boost in BC3S1, corresponding to a linkage drag reduction by a factor 2, to be compared to the factor 10 found previously when the target locus is in a cold region. Moreover, we also wondered whether using HR instead of Boost (higher increase of recombination in hot regions compared to Boost, Fig. 1) might be advantageous. We find that there is little interest in using HR: the segment length is 0.02 Mb under HR instead of 0.03 Mb under Boost in BC3S1. Thus in a region where recombination is already significantly present, the initial level of recombination is sufficient to have a high decrease of the linkage drag and so neither Boost nor HR are very advantageous.

### Effect of increased recombination on the recovery of the recurrent genome

#### For a reference situation

We followed the recovery of the recurrent genome *via* the (weighted) proportion of donor alleles remaining in the genome (see Materials and Methods). Graphical genotypes are displayed in Fig. 3, providing insight into the situation, while Fig. 4.B shows the mean proportion at each generation. In the absence of selection on the proportion of donor genome (*i.e.,* no background selection), we expect the proportion of donor alleles to decrease by half at each generation, and this is what we observe up to BC2 (proportion of the donor genome: 100% in F0, 50% in F1, 25% in BC1 and 12.5% in BC2), for both levels of recombination (Fig. S3.B). However, once background selection is implemented (from BC3 onward), the donor proportion decreases faster than in the absence of selection: in BC3, 1.6% of the genome is donor under WT and 2.1% under Boost. At the end of the selection program, the proportion of donor genome is lower under Boost than under WT. Specifically, there remains about twice as much donor genome under WT compared to Boost (Fig. 4.C): when one reaches BC3S1, on average, 2.3% of the genome comes from the donor under WT while the proportion is 0.9% under Boost. Note that it is possible for the proportion of donor genome to *increase* after one round of selfing as some markers that were heterozygous will become homozygous for the donor allele. Overall, under Boost, the selection against the donor alleles is more effective than under WT during the round of selfing: while the proportion of donor genome increases under WT, it continues to decrease under Boost, ending with a lower proportion under Boost in BC3S1.

Considering now the situation where the target locus is in a region that is relatively hot for recombination in WT (Fig. 1), we find that WT is better than having increased recombination (Fig. S5.B), be it HR or Boost although HR is slightly better compared to Boost: in BC3S1, on average 0.07% of the genome is donor in WT, 0.46% in HR and 0.57% in Boost.

Although the previous section showed that increased recombination has benefits for decreasing linkage drag, it simultaneously hampers the recovery to the recurrent genome on the rest of the genome as seen in Fig. 3 and Fig. 4. This is true both for the chromosome carrying the target and for the other chromosomes. As illustrated in Fig. 3 and Fig. 4.D, in WT, most of the remaining donor genome is carried by the segment around the target locus (in BC3S1, 96% of the remaining donor genome is due to linkage drag in WT against 36% in Boost). Furthermore, Boost leads to many small segments of donor genome, distributed throughout the different chromosomes. These segments are difficult to remove, an intrinsic drawback of the increased recombination approach. Nevertheless, as we have seen before, adding one more generation of backcross strongly cleans up the background, in particular under Boost: for instance in BC4S1, the percentage of the donor genome due to linkage drag is then nearly 100% in WT and 95% in Boost.

#### General trends of the advantage of increased recombination

As a first aspect, consider adding additional generations of backcross. We find that this is an effective way to get rid of the donor genome even further. For instance, when the program goes up to BC4S1, the difference between the two levels of recombination becomes even more pronounced, with nearly ten times more donor genome remaining under WT compared to Boost: in BC4, 1% of the genome is donor under WT (1.8% in BC4S1) while it is 0.2% under Boost (0.2% in BC4S1). The advantage of Boost increases when adding further generations but at a diminishing rate. For instance the ratio representing the mean proportion of donor genome under WT divided by the one under Boost is 5.3 in BC4, 11 in BC5, 13 in BC6, 15 in BC7 and 17 in BC8 (Fig. S4.C). This is not surprising given that nearly all of the donor genome is coming from the linkage drag starting from BC4 in WT and starting from BC6 under Boost (Fig. S4.D).

When changing other parameter choices in the backcross selection program we see similar trends to the ones that were found in the case of linkage drag. For instance, increasing the population size results in better recovery of the recurrent genome (Fig. S3.B and C): in BC3S1, on average 2.9% of the genome is donor under WT and 1.2% under Boost for 200 individuals, while the averages are 1.5% under WT and 0.5% under Boost for 1000 individuals.

However the trends are not always advantageous for increased recombination. At least that is the result specifically when the target locus is already in a relatively hot region in WT: there we find that it is *disadvantageous* to use increased recombination (be it HR or Boost) as it increases the difficulty of cleaning up the genetic background: in BC3S1, 15% of the remaining donor genome is in the background in WT, 85% under Boost and 88% under HR.

### Selecting for the second best plant instead of the best one

All of the previous results were obtained supposing that it is possible to select the best individual at each generation. In the case that the best individual is not available (it could accidently die or give rise to no seeds), we looked at the effect of selecting the *second* best plant at each generation (Fig. S6). What we observe is similar to when the best individual is selected, although not as effective for both the reduction of the linkage drag and the recovery of the recurrent genome. Specifically, when the second best plant is selected, in BC3S1, the average donor segment size is 7.4 Mb in WT and 1.8 Mb in Boost (to be compared to 5.2 Mb and 0.5 Mb under WT and Boost respectively when the best individual is selected). Similarly, in those conditions the average proportion of donor genome is 3.27% under WT and 1.75% under Boost, to be compared to 2.28% and 0.85% under WT and Boost, respectively, when the best individual is selected. We find that under WT, the clean-up of the background is similar whether the best or second best plant is kept (at BC3S1, 96% of the remaining donor genome is due to the donor segment around the target locus when the best is selected and it is 95% when the second best is kept). On the contrary, under Boost, selecting the second best results in a lower relative contribution of the background: the donor segment around the target locus represents 36% of the remaining donor genome when selecting the best individual while it represents 42% when the second best is kept.

## DISCUSSION

When increasing the recombination level, we find that it is most useful to do so in the regions that are initially poor in crossovers.

Specifically, Boost is advantageous when the target locus is in a region that is cold in WT (e.g. in pericentromeric regions): in WT, the decrease of the linkage drag is restricted inthe zone without recombination while most of that zone recombines under Boost, allowing for a sharp decrease of the linkage drag during the foreground selection. In this situation, Boost is also globally beneficial, even more so when one more generation is added (BC4S1 instead of BC3S1), allowing for a quite efficient clean-up of the genetic background. On the contrary, when the target locus is in a hot region, WT is sufficient to drastically decrease the linkage drag during foreground selection. WT is also better globally because under increased recombination many small donor blocks are produced in the genetic background and that is disadvantageous during background selection (Hillel *et al.* 1990). Thus in situations of target loci lying within hot regions, adding further recombination is generally detrimental. To be explicit, although it does decrease linkage drag (at the end of the foreground selection, in our simulations the mean donor segment size is 0.07 Mb under WT, 0.03 under Boost and 0.02 under HR), that is not enough to balance the associated pollution of the genetic background by the donor genome. By increasing the population size, it is possible to get crossovers closer to the target locus (Hospital *et al.* 1992, Frisch *et al.* 1999a), even under WT, although it is never better than when using Boost, in the range of population sizes studied here. Let us note nevertheless that under Boost, the change in the population size has a smaller effect compared to WT. Despite this, Boost is still better overall.

It is also possible to decrease the linkage drag in WT by adding more generations. At some point (around BC5), almost all of the remaining donor genome will be due to the segment around the target locus. Thus, selecting to decrease the proportion of donor genome will in fact select to decrease the size of the donor segment around the target locus. This is particularly strong under WT, as the segment around the target locus is large and there is nearly no unwanted donor genome in the genetic background. This property allows one to justify the importance of foreground selection: in a long-term backcross selection program, the amount of residual donor genome will be nearly completely determined by the quality of the foreground selection.

As an outlook towards potential impact on real breeding programs, one may ask whether increased recombination could speed-up backcross programs by reducing the number of generations used. Focusing on the main selection scheme of our work, namely foreground selection for two generations followed by one or two background selections followed by selfing, we see in Fig. 4B that BC3S1 when using Boost is better than BC4S1 under WT (which of course is itself better than BC3S1 under WT). Thus, under certain conditions, increased recombination may speed-up backcross programs while simultaneously improving performance. The advantage of increased recombination is even more striking when considering the long backcross programs displayed in Fig.S4C where Boost at BC4 outperforms all generations of WT including BC8. There may of course be other strategies for shortening generation times thanks to increased recombination. For instance one may attempt to apply foreground selection for a single generation; this would correspond to selecting double recombinants (crossovers on both sides of the target locus) in BC1. Clearly the performance of such an approach will depend a lot on how the recombination landscape around the target locus is changed when going from WT to Boost but it may provide a practical way to speed-up backcrossing programs for introgressing segments lying in certain cold regions.

## DECLARATIONS

### Ethics approval

Not applicable

### Consent to participate

Not applicable

### Consent for publication

Not applicable

### Availability of data and material

The parameters files for each simulation and the analysis scripts are included in the R package containing the simulation scripts.

### Code availability

The scripts for the simulations can be downloaded as an R package on https://sourcesup.renater.fr/frs/download.php/latestfile/2217/carebBC.tar.gz

### Authors’ contributions

MF and OCM conceived, designed and supervised the study. ET performed the simulations. ET, MF and OCM performed the analysis and interpreted the results. ET, MF and OCM wrote and edited the manuscript. All authors read and approved the final manuscript.

## Acknowledgements

We thank S. Mezmouk, E. Jenczewski, J.C. Motamayor, D. Livingston, C.E. Durel and R. Mercier for constructive comments and discussions and A. M. Chèvre for her comments on the manuscript.

---

**Figure S1** Recombination landscape for the 10 chromosomes of *Brassica rapa.* The WT landscape is represented in black, HR is in green and Boost is in red. The solid lines represent the profiles obtained from experimental data (WT and Boost, data from Pelé *et al.* 2017) while the dotted line is the simulated profile (HR). The centromere positions are represented using blue bars (Mason *et al.* 2016).

**Figure S2** Recombination landscape for the 12 chromosomes of *Oryza sativa*. The WT landscape is represented in black while the HR is in green; both are obtained experimentally (Mieulet *et al.* 2018). The centromere positions are represented by blue bars (Mizuno *et al.* 2018).

This figure shows the effect of HR recombination on the shape of the recombination landscape, which is reproduced by simulation for *B. rapa*.

**Figure S3** (A) Mean size of the heterozygous segment around the target locus, in Mb, as a function of the generations, in *B.rapa*. (B) Mean proportion of the donor genome, in percentage, as a function of the generations. The insert represents a zoom on the last generations, from BC2 onwards. (C) Ratio of the mean proportion of donor genome under WT over the proportion under Boost as a function of the generations. (D) Mean proportion of the remaining donor genome that is due to the linkage drag, calculated as the part of the remaining donor genome that is due to the heterozygous segment around the target locus, as a function of the generations. The measures under WT are represented in black while those under Boosted recombination are in red. Different population sizes are shown: 200 plants per generation in dashed lines, 400 in solid lines and 1000 in dotted lines. In the situations represented in this figure, the target locus is in a cold region, and the selection scheme goes up to BC3S1. The best individual is kept at each generation, following the selection criterion appropriate for each generation (BC1 and BC2: foreground selection, and thereafter background selection).

**Figure S4** (A) Mean size of the heterozygous segment around the target locus, in Mb, as a function of the generations, in *B.rapa*. (B) Mean proportion of the donor genome, in percentage, as a function of the generations. The insert represents a zoom on the last generations, from BC2 onwards. (C) Ratio of the mean proportion of donor genome under WT over the proportion under Boost as a function of the generations. (D) Mean proportion of the remaining donor genome that is due to the linkage drag, calculated as the part of the remaining donor genome that is due to the heterozygous segment around the target locus, as a function of the generations. The measures under WT are represented in black while those under Boosted recombination are in red. The selection scheme goes up to different number of generations: BC3S1 in solid lines, BC4S1 in dashed lines and BC8 in dotted lines.

In the situations represented in this figure, the target locus is in a cold region and there are 400 plants per generation. The best individual is kept at each generation, following the selection criterion appropriate for each generation (BC1 and BC2: foreground selection, and thereafter background selection).

**Figure S5** (A) Mean size of the heterozygous segment around the target locus, in Mb, as a function of the generations, in *B. rapa*. (B) Mean proportion of the donor genome, in percentage, as a function of the generations. The insert represents a zoom on the last generations, from BC2 onwards. (C) Ratio of the mean proportion of donor genome under WT over the proportion under Boost as a function of the generations. (D) Mean proportion of the remaining donor genome that is due to the linkage drag, calculated as the part of the remaining donor genome that is due to the heterozygous segment around the target locus, as a function of the generations.

The measures under WT are represented in black while those under Boosted recombination are in red, and those under HR are in green. The target locus is either in a cold region (solid lines) or in a hot region (dashed lines). HR was not considered when the QTL was in a cold region as the effect of increased recombination mainly influenced the linkage drag and HR was not different from WT in the cold region.

In the situations represented in this figure, there are 400 plants per generation and the selection scheme goes up to BC3S1. The best individual is kept at each generation, following the selection criterion appropriate for each generation (BC1 and BC2: foreground selection, and thereafter background selection).

**Figure S6** (A) Mean size of the heterozygous segment around the target locus, in Mb, as a function of the generations, in *B.rapa*. (B) Mean proportion of the donor genome, in percentage, as a function of the generations. The insert represents a zoom on the last generations, from BC2 onwards. (C) Ratio of the mean proportion of donor genome under WT over the proportion under Boost as a function of the generations. (D) Mean proportion of the remaining donor genome that is due to the linkage drag, calculated as the part of the remaining donor genome that is due to the heterozygous segment around the target locus, as a function of the generations.

The measures under WT are represented in black while those under Boosted recombination are in red. Either the best (solid lines) or the second best (dashed lines) plant is kept at each generation.

In the situations represented in this figure, the target locus is in a cold region, there are 400 plants per generations and the selection scheme goes up to BC3S1.

