## Supplementary figures and images for "Enhancing backcross programs through increased recombination"

### Supplementary figures 1 to 6

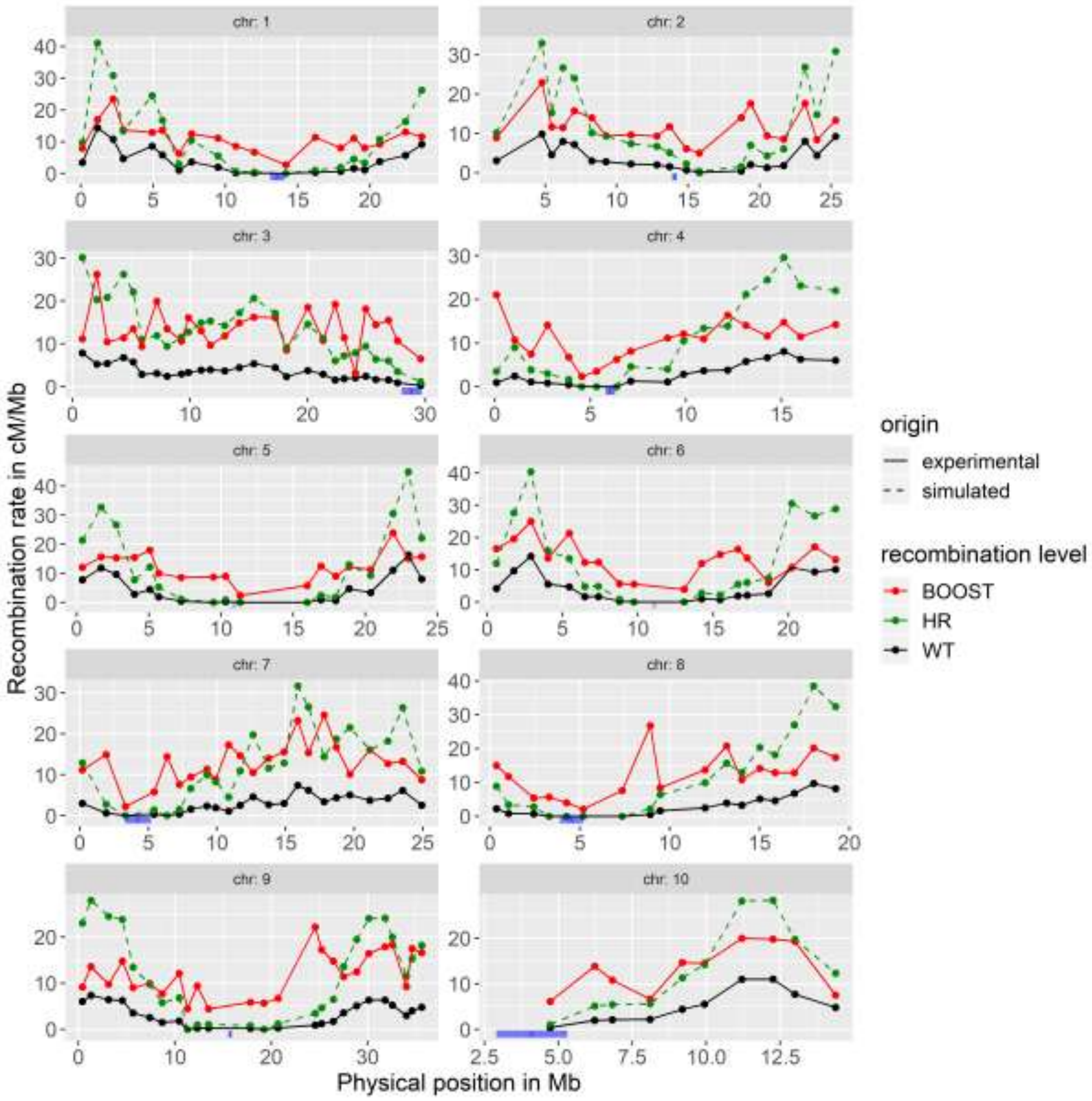


**Figure S1**

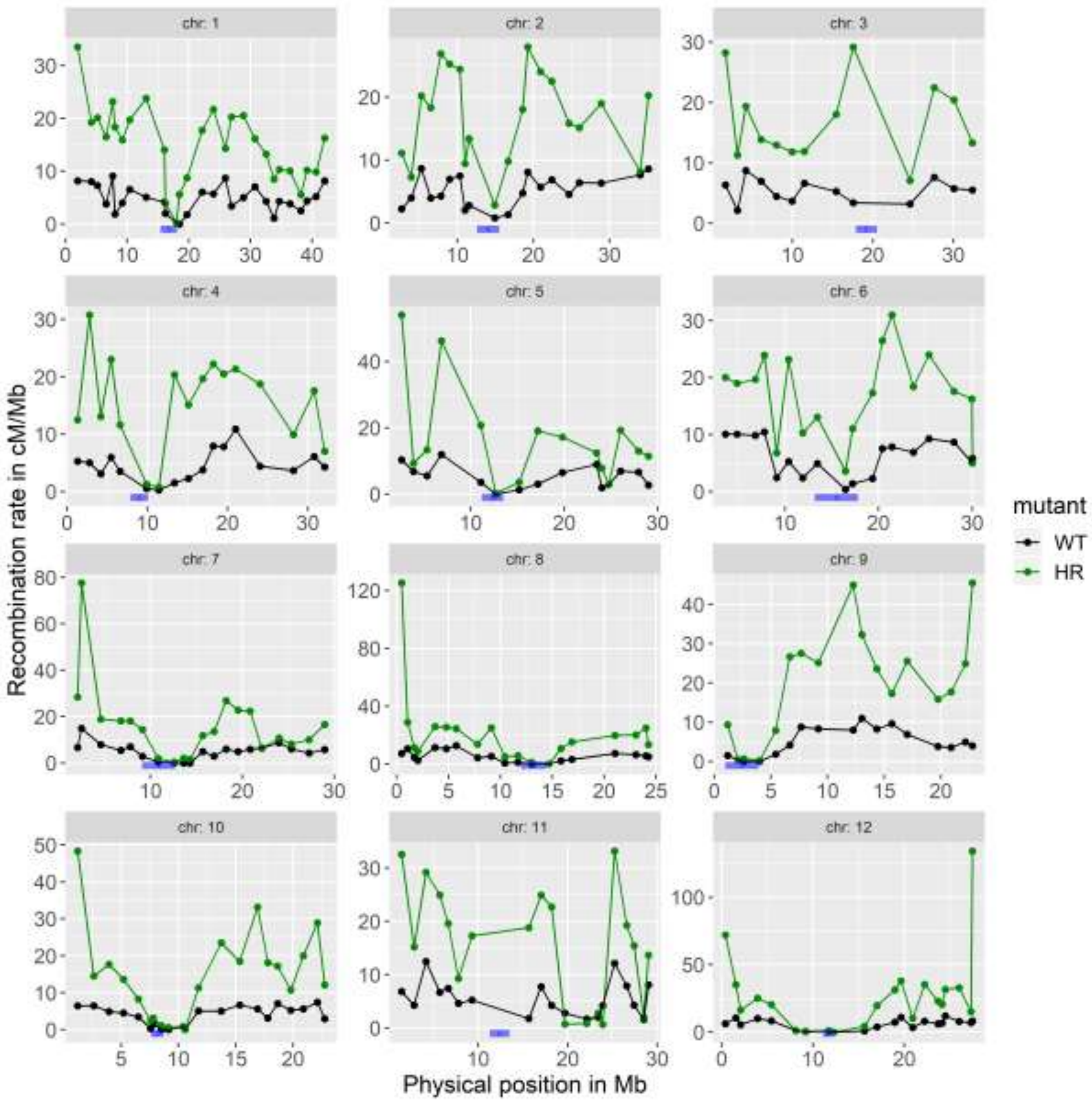

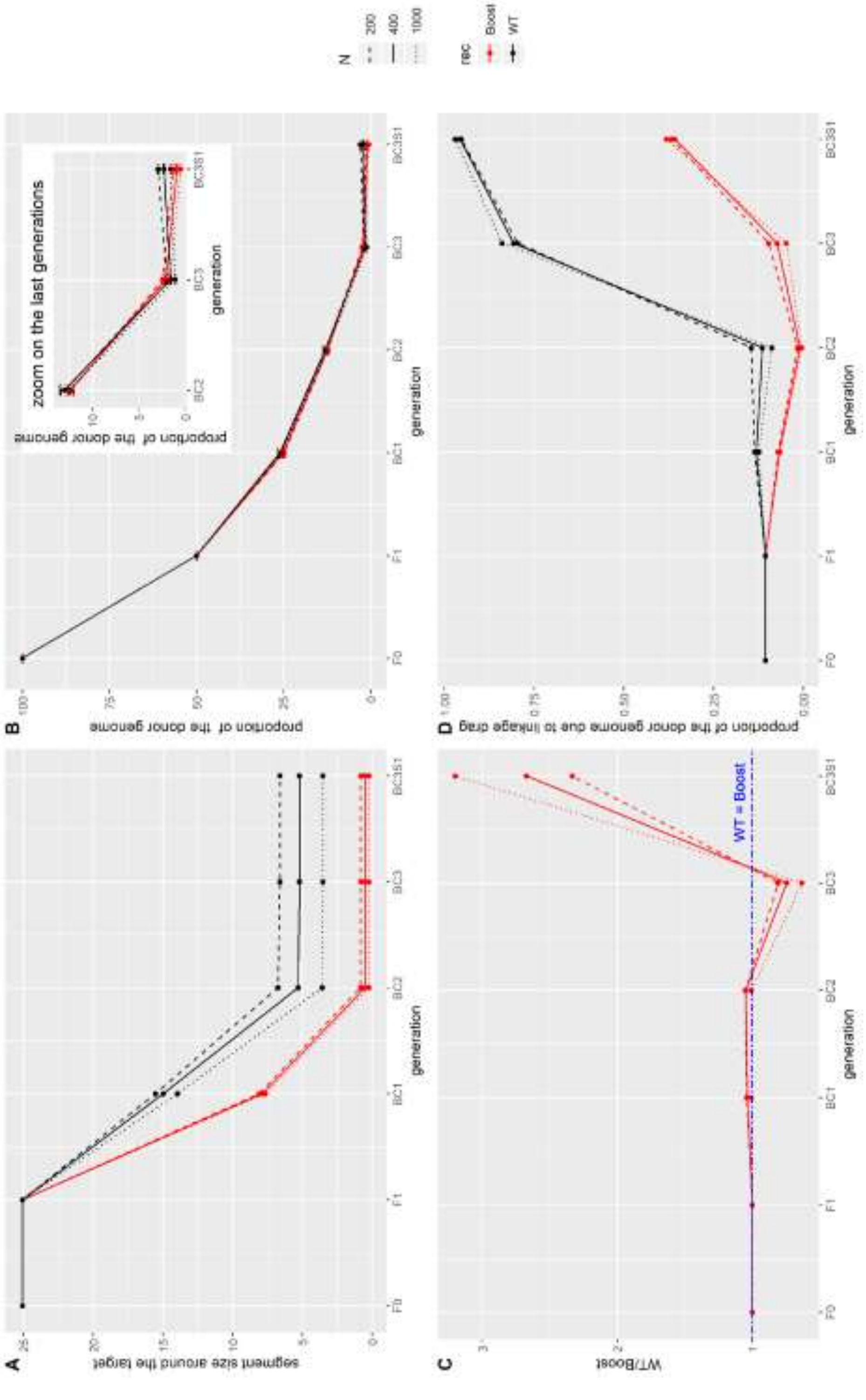

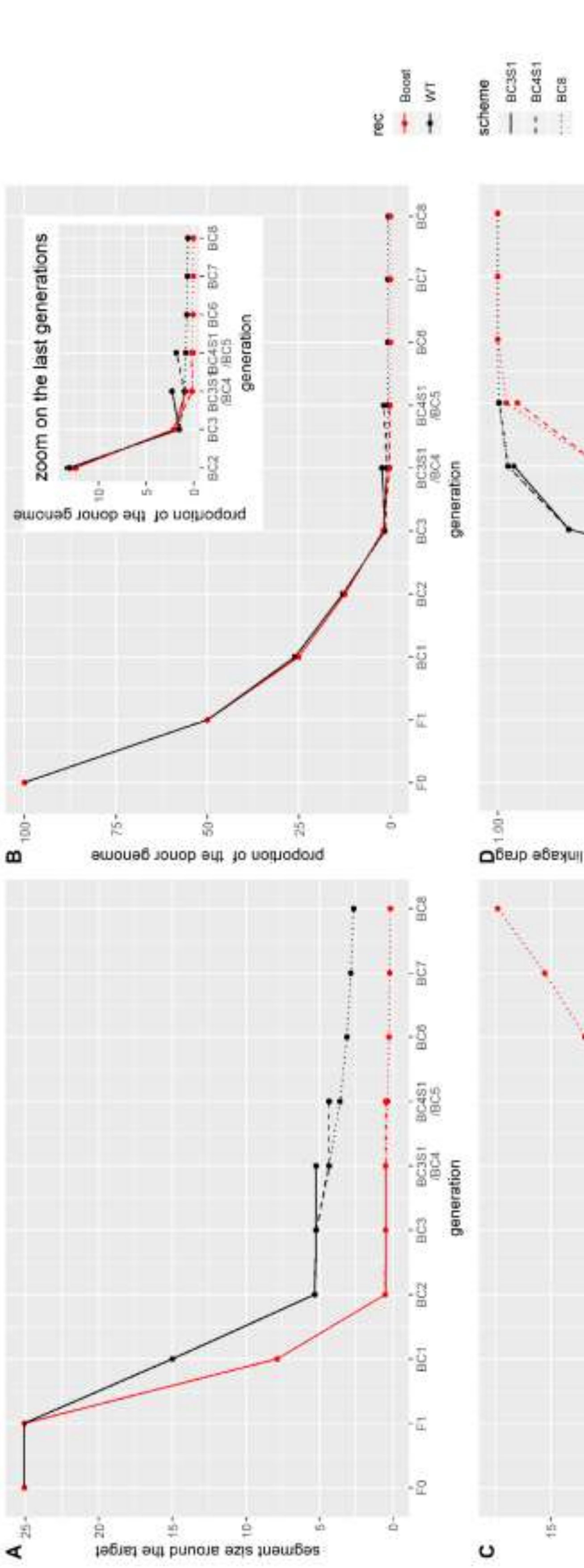

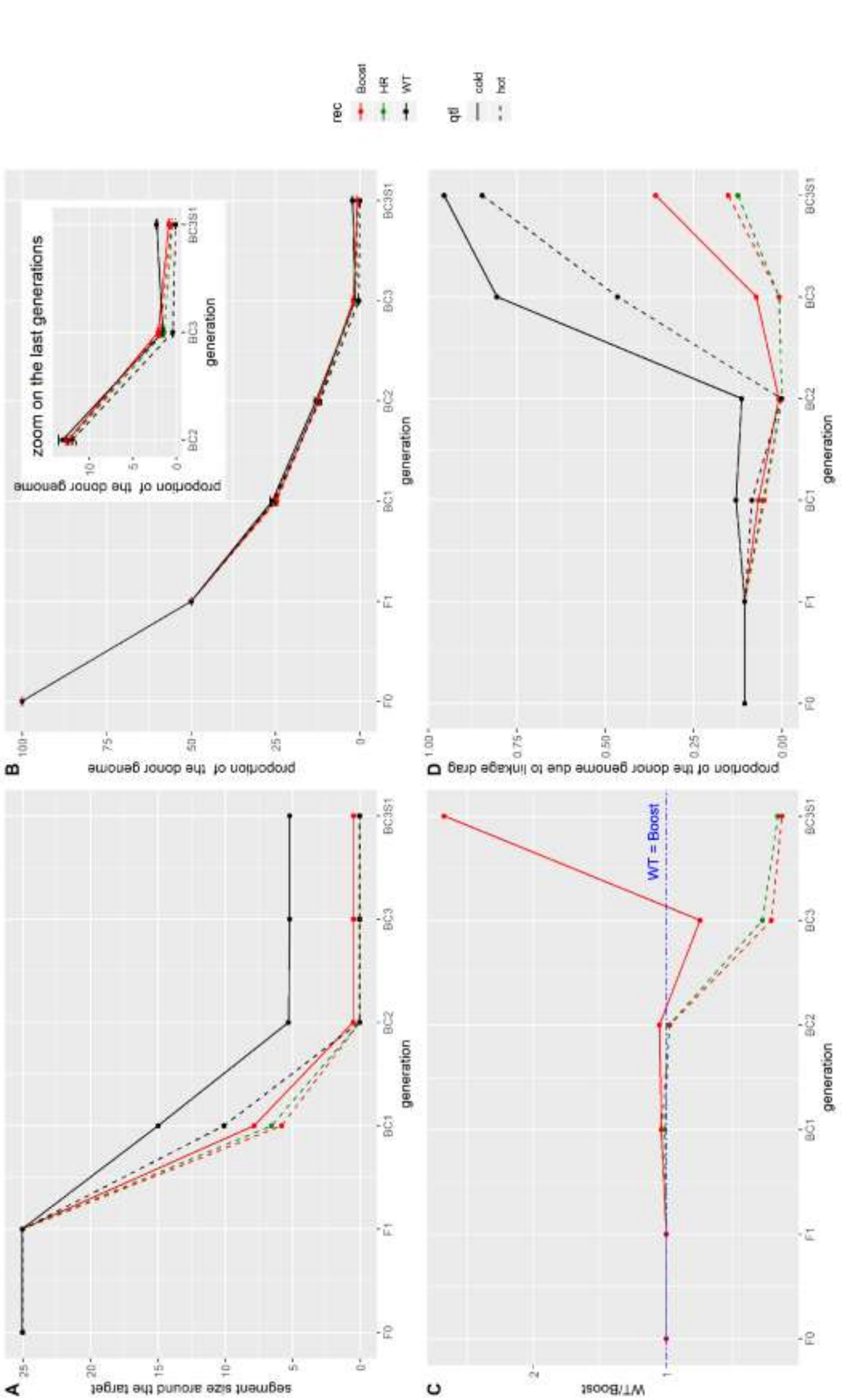

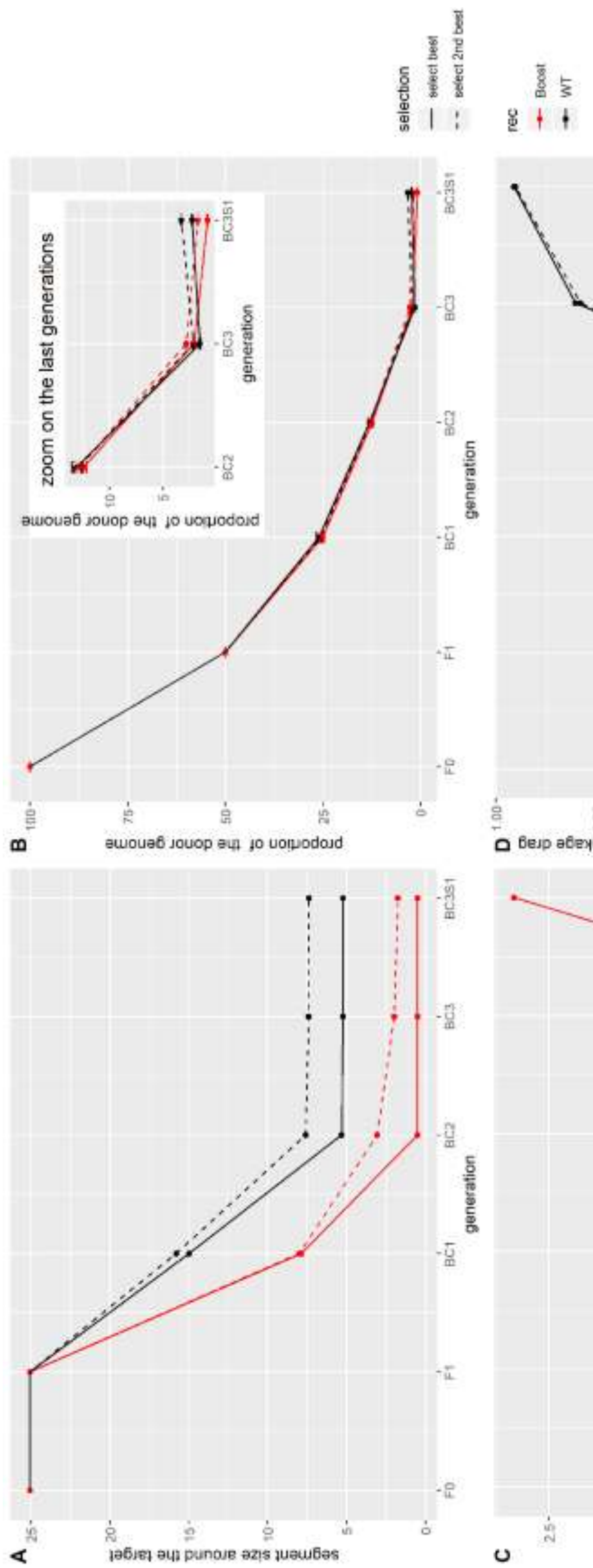
